# Fragmented Spatial Maps from Surprisal: State Abstraction and Efficient Planning

**DOI:** 10.1101/2021.10.29.466499

**Authors:** Mirko Klukas, Sugandha Sharma, YiLun Du, Tomas Lozano-Perez, Leslie Kaelbling, Ila Fiete

**Affiliations:** BCS & McGovern Institute, MIT, Cambridge MA 02139, USA; EECS & CSAIL, MIT, Cambridge MA 02139, USA

## Abstract

When animals explore spatial environments, their representations often fragment into multiple maps. What determines these map fragmentations, and can we predict where they will occur with simple principles? We pose the problem of fragmentation of an environment as one of (online) spatial clustering. Taking inspiration from the notion of a *contiguous region* in robotics, we develop a theory in which fragmentation decisions are driven by surprisal. When this criterion is implemented with boundary, grid, and place cells in various environments, it produces map fragmentations from the first exploration of each space. Augmented with a long-term spatial memory and a rule similar to the distance-dependent Chinese Restaurant Process for selecting among relevant memories, the theory predicts the reuse of map fragments in environments with repeating substructures. Our model provides a simple rule for generating spatial state abstractions and predicts map fragmentations observed in electrophysiological recordings. It further predicts that there should be “fragmentation decision” or “fracture” cells, which in multicompartment environments could be called “doorway” cells. Finally, we show that the resulting abstractions can lead to large (orders of magnitude) improvements in the ability to plan and navigate through complex environments.

## Introduction

Contextual reorientation [1], in which behavior, state estimates, or meaning are suddenly reevaluated and reanchored based on new or altered contextual information from the world, are universal phenomena in psychology. One interesting set of examples is the parsing of garden-path sentences such as “Time flies like an arrow, fruit flies like a banana” or “The woman brought the sandwich from the kitchen tripped” [2]. In the latter there is a sudden reorientation upon hearing the word tripped, so that *the woman* becomes the person *who was* brought the sandwich rather than the person bringing the sandwich. Similarly, spatial reorientation and reanchoring can occur when entering a building lobby from the outside or entering a different looking room from another one. Such reanchoring or reorientation events may constitute the basis on which we segment the continuous stream of experience into episodes or chunks to structure experience and memory [3–5].

In the brain, grid cells construct continuous 2-dimensional Euclidean maps of small environments [6] by the integration of self-movement cues as the animal explores the space, Fig. 1a. The advantage of such velocity integration-based Euclidean representations is that they provide a consistent encoding of seen and unseen locations and independent of paths taken to get there, making it possible to compute novel shortcut paths and perform spatial inference between locations [7–12].

**Figure 1:**
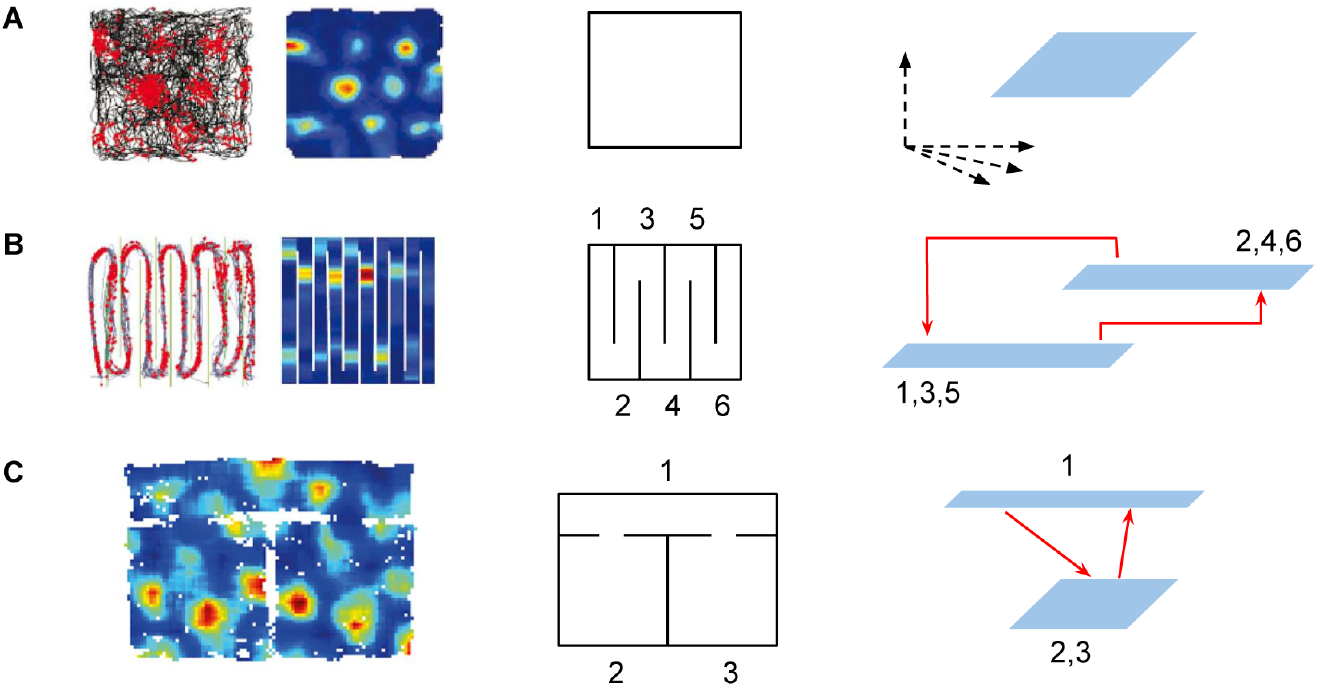
Map fragmentation in MEC. **A-C:** *Left,Middle:* Firing fields of grid cells in various environments (firing fields in A,B from [13] and in C from [14]). Environments are illustrated schematically in the middle column (open field, hairpin, and two-room with hallway). *Right:* Schematic map fragmentations of the environments. Blue regions are submaps, or regions with a continuous representation in the state space of the multi-module grid cell population. Solid arrows indicate discontinuous jumps in grid phase, which we interpret as transitions between submaps.

However, in more complex spaces with compartments or subregions [13, 14], Fig. 1b-c, the grid cell spatial representation is often not a globally consistent 2-dimensional map that can be predicted by path integration even when the agent moves smoothly and continuously across the space. Instead, the neural representations exhibit a fractured structure across the space, referred to in the literature as map fragmentation. We hypothesize a direct connection between cognitive reanchoring and the formation of fragmented spatial maps including the general phenomenon of remapping.

It is well known that between different environments, place and grid cells “remap”: Representations of different environments involve different (if overlapping) sets of place cells and the spatial relationships between place cells in one environment are not preserved in the other [15]. Grid remapping is seemingly more subtle, but consistent with place cell remapping: within modules, grid cells maintain their relationships and thus shift their phases coherently, but across modules, there are differential or incoherent shifts in phase [16]. Remapping is typically studied by discontinuously transplanting subjects from one environment to another or by switching non-spatial cues [15, 17]: by inducing large spatial changes through a journey in a closed container or vehicle or cueless corridor where the subject cannot easily determine its spatial displacement, or by altering olfactory or visual cues within the same space. Map fragmentation – because it involves the continuous movement of an agent through a stationary environment – is viewed differently from remapping, in which the emphasis on an externally-induced environmental change [15, 18]. However, map fragmentation should be viewed as remapping or reanchoring within an environment; here we seek to provide a unified model of map fragmentation and explore its potential utility.

A cost of map fragmentation is the loss of the ability to perform path-integration based computations across the environment [7–11]. We hypothesize that map fragmentation is a solution to multiple problems: First, it solves the problem of the accumulation of path integration errors that prevent the formation of consistent maps over larger spaces, resulting in the formation of smaller but consistent Euclidean maps. Thus, map fragmentation enables spatial inference and shortcut behaviors within each submap. Second, each submap represents a state abstraction in which contiguous locations are clustered together, and combining these abstractions with links between them can permit efficient and hierarchical representation and planning. Third, submaps can combine more globally to form a “topometric” map, a representation with enough expressiveness for topologically non-trivial cognitive spaces beyond real space, that preserves the advantages of both local metric structure and global hierarchy and abstraction.

Here, we propose a simple online rule for map fragmentation that avoids the large memory, time complexity and data-inefficiency of offline algorithms, and show that the resulting rule is a good potential model of map segmentations observed in grid and place cells. Finally, we demonstrate by implementing efficient random tree search algorithms that map fragmentation can facilitate efficient planning relative to using global maps, leading to a massive speed-up in complex and large environments without repeated substructures.

## Results

### Map fragmentation as clustering: an offline baseline

We propose that remapping across environments and fragmentation within environments can be considered to be a clustering problem: At each sampled location, the question is whether it should be categorized as a part of the most recently used map, or be assigned to a different one. A sensible answer would be that sufficiently “similar” locations should be assigned to the same map (cluster), while sufficiently different ones should be assigned to different maps (clusters).

We view a map as a (local) world model that enables the prediction of sensory inputs at any location within the map. Thus, we consider that a key metric for map fragmentation may be predictability or surprisal, Fig. 2. A similar metric has been used in robotics methods for simultaneous localization and mapping (SLAM) [19]. Specifically, sets of poses (locations and orientations) where the predictability of external observations remains high while moving between them (“contiguous regions”) should be clustered together into one map, Fig. 2. This view complements the use of other metrics that have been implemented in offline settings to construct spatial maps, including the graph Laplacian [20] and successor representation [21] methods, both of which use temporal proximity as their metric (indeed, under a random exploration policy, the successor representation is closely related to the graph Laplacian). Our primary focus here is on how biological and artificial agents might generate sensible maps in an online fashion. Secondarily, we use the metric of prediction or surprisal to generate these online fragmentations. In Discussion, we will consider how additional metrics can be used within the same online framework.

**Figure 2:**
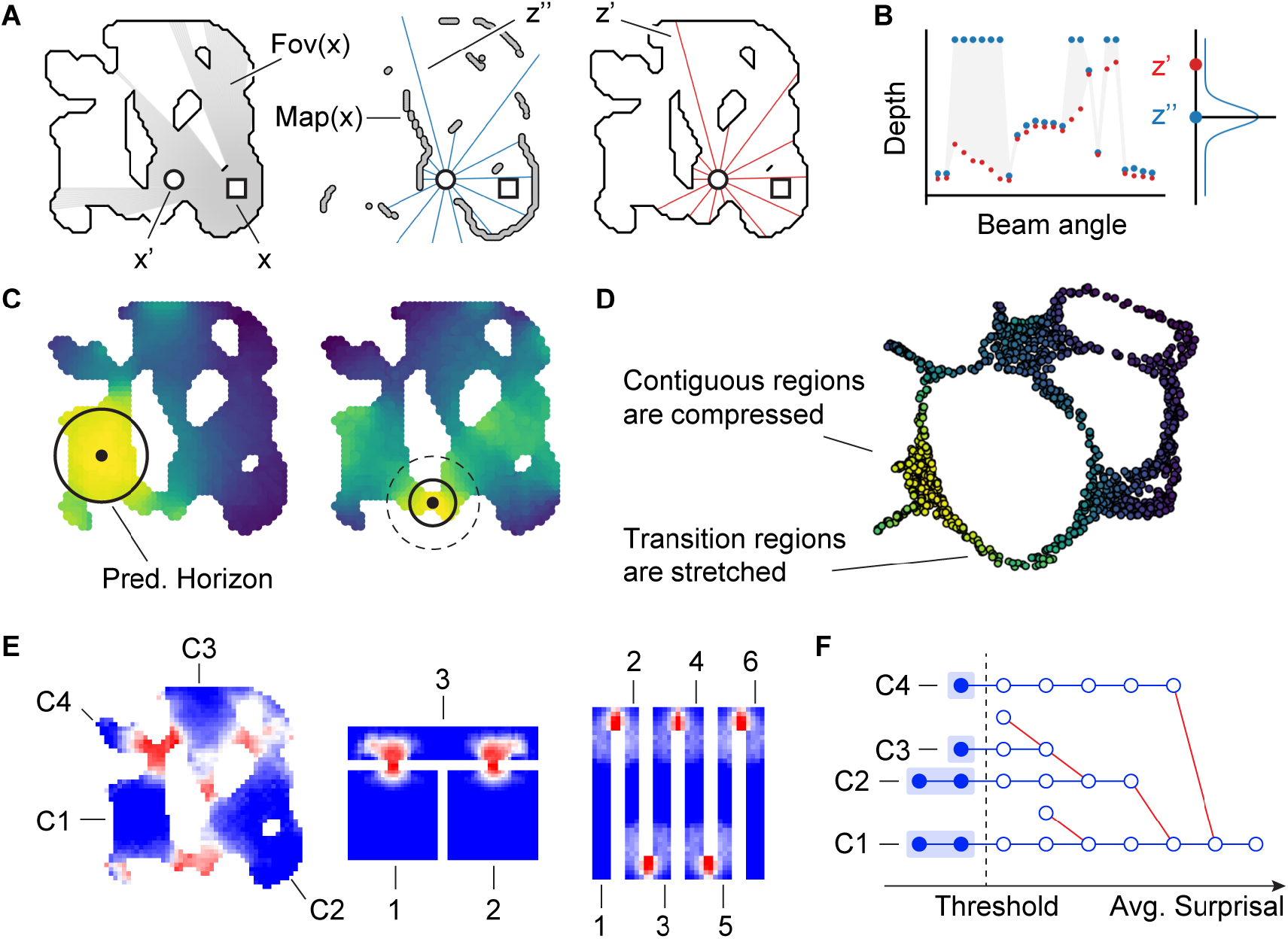
Fragmentations from predictability-based clustering. **A:** *Left*. Two agents (circle and square) in an organically shaped environment, with gray indicating the square agent’s field of view. *Middle*. A set of range sensor observations (blue, *z’’*) by the circular agent, shown on the map built from the square-agent’s observations. *Right*. A set of range sensor observations (red, *z’*) by the circular agent, shown on the actual environment. **B:** The blue observations in A constitute the square agent’s prediction of the circular agent’s measurements, with a prediction model given by a multivariate diagonal Gaussian with means given by the blue measurements, and which is evaluated at the vector of actual measurements in red. Greater vertical deviations between red and blue dots correspond to larger prediction errors. **C:** Each black dot represents a fixed reference location. All locations (pixels) in the map are colored by their predictability from the reference location. Solid black circle: the prediction horizon at that reference location: The horizon is large in open (contiguous) spaces, and small at bottleneck (transition) regions. **D:** An Isomap embedding of the environment based on mutual predictability gives rise to a warped embedding: distances are large when predictability is low. **E:** Locations in three environments, colored by their *average surprisal*. Numbered subregions correspond to connected components with average surprisal below a threshold level. **F:** A hierarchical clustering tree for the evolution of connected subregions for the first environment from E: Although the delineation of subregions depends on the choice of threshold, some subregions are relatively persistent and thus robust, maintaining their identity over a range of thresholds.

Define a model 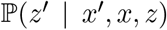 that predicts the sensory input *z’* at pose *x’*, based on the sensory input *z* at pose *x* (Fig. 2a,b; see Methods for details). The sensory observations and their predictions are given in terms of a range sensor centered on the agent, in the actual environment (Fig. 2a, right) or in a reconstructed map based on the observations *z* (Fig. 2a, middle), respectively. For each pose *x*, we delineate the surrounding region where predictability remains above threshold; this, by definition, is a contiguous region. We call the boundary of the region the prediction horizon for *x*. The radius of the prediction horizon varies depending on location within the environment, Fig. 2c. We can use the mutual surprise between poses, which we define as 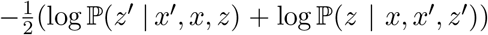 (see Methods for details), as a measure of proximity that we illustrate with an Isomap embedding [22] of the environment (Fig. 2d). In this visualization, contiguous (high predictability) regions are compressed, while transition or bottleneck regions (low predictability) are stretched.

Finally, we define the average surprisal (see Methods) of a pose x by averaging over the mutual surprise of all nearby poses at a fixed Euclidean distance, and apply a clustering procedure similar to DBSCAN [23]^1^. The procedure computes the connected components of all locations whose average surprise lies below a fixed threshold and decomposes the map into core fragments and transition regions, Fig. 2e. Additionally, in order to make an informed choice about the fragmentation threshold, we compute a contour tree (cf. [24]) of the surprisal values, which provides a visualization of how the connectivity of space evolves with increasing thresholds, Fig. 2f. As we see, there are regions of the contour tree that are relatively robust to the detailed threshold choice, providing similar connectivities over a range of threshold values.

The surprisal-based segmentations align well with both intuitive fragmentations and with neural data (cf. Fig. 1), suggesting that predictability may be a key and principled objective for map segmentation decisions.

However, the algorithm is offline, requiring full exploration of the space before it can generate the fragmented map. This is unlike in experiments, where animals generate map fragmentations in real-time as they explore an environment [14]; in non-spatial contexts too, there is evidence that event boundaries are defined in real-time [5, 25]. The algorithm also has high complexity, requiring fine spatial discretization and a large memory and computational buffer for the storage of and computation on the full predictivity matrix over all pairs of positions in the space. The same is true for Laplacian and successor matrix-based methods. Further, there is an additional gap between observed map fragmentations in biology and the latter two algorithms because while they provide multi-scale representations of the space (in the form of eigenvectors of some similarity matrix), they are not actually fragmentations of the environment, Fig. S3,S4.

### Online fragmentation based on predictability: Our model

We next build a simple and biologically plausible online map fragmentation model based on short-term prediction error as an efficient proxy for surprisal, with the goal of generating fragmentations that are consistent with the principled offline clustering-based algorithm above. Our model is an agent that integrates its velocity as it explores an environment to update its pose estimate, and uses a short-term memory (STM) and a long-term memory (LTM) to make predictions about what it expects to see next.

The sensory observations for the online model consist of the activities of a population of cells that encode the presence of environmental boundaries at some distance, similar to boundary vector cells (BVCs) [26, 27] in entorhinal cortex or boundary-coding cells in the occipital place area [28]. These encode a binary, idiothetically-centered local view of the space^2^ (with observation field-of-view angle *ϕ*), Fig. 3a,c. The velocity-based position estimates are represented by a population of idealized grid cells from multiple modules. For simplicity and to match the experimental setups in [13, 14] we assume that the pose angle is specified by a global orienting cue – effectively, the agent has access to its true head direction. The STM consists of an exponentially decaying moving average of recent observations, each shifted according to the internal velocity estimate of the agent, Fig. 3d. The STM is used to generate the prediction for the next observation (motivated by [19]), and a normalized dot product between the prediction and the current observation (BVC activity) yields our predictability signal (Fig. 3b,e). Due to its implementation as a moving average, STM activity slightly lags BVC activity. While high predictability is maintained along a trajectory, no fragmentation occurs. Once the predictability signal dips below a threshold, then at the first subsequent stabilization of spatial information, signaled by predictability returning to threshold, a fragmentation event is triggered (Fig. 3b). The fragmentation decision thus segments or chunks the continuous stream of experience based on the content of the experience, with similar observations chunked together and separately from dissimilar ones. It is an event that is discrete in space and time and drives a discrete fracture or fragmentation in the in-
ternal map that translates to a discontinuous jump in the large encoding space of the multi-periodic grid cell population.

**Figure 3:**
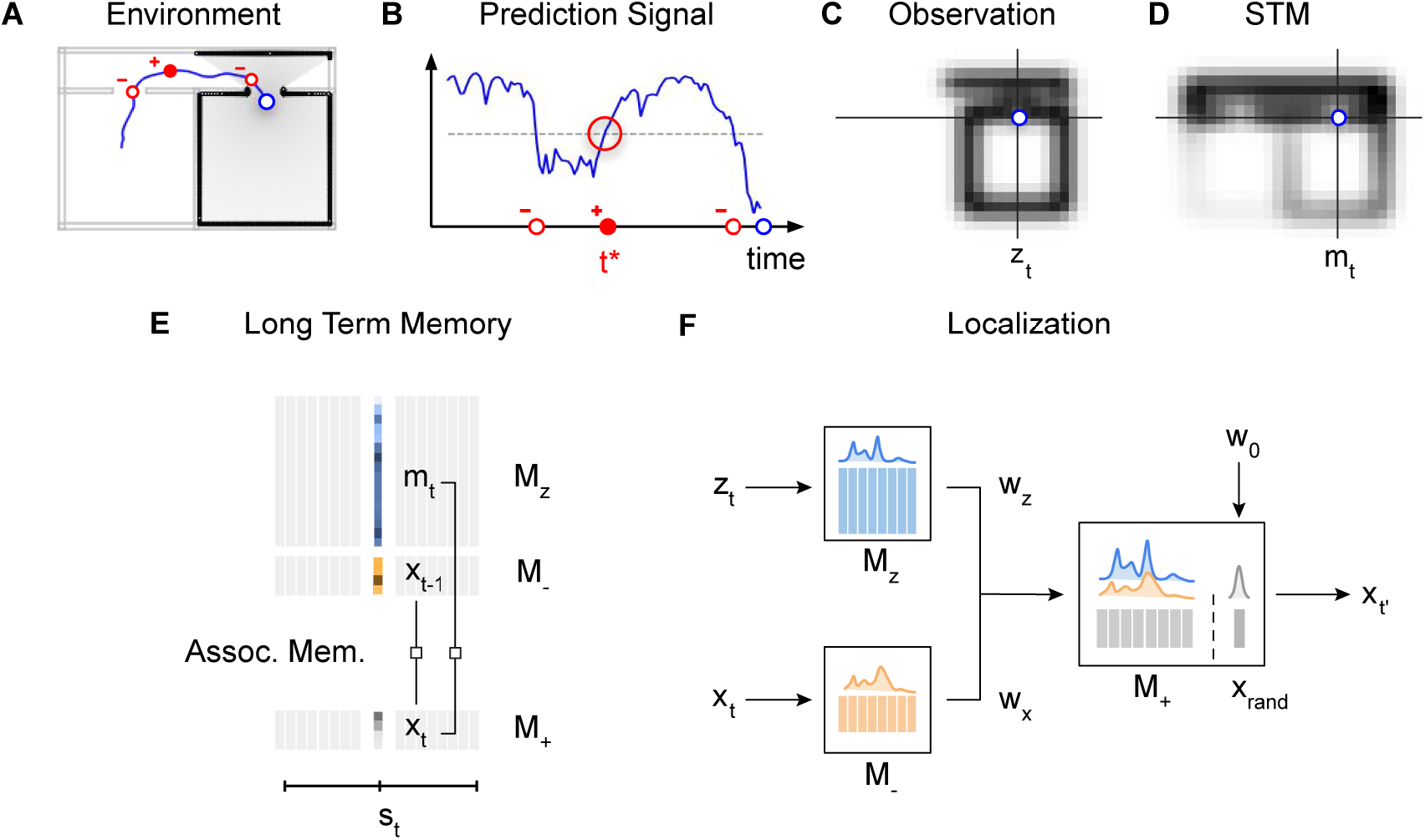
Online fragmentation model. **A:** Sample trajectory (blue) through a “2-room” environment highlighting three distinct events (red circles) that map to the events in B. Gray area: the agent’s field of view at the agent’s current location (blue circle). **B:** Prediction signal for the trajectory in A. Red dots indicate when the prediction crosses a threshold (dashed line), with the filled dot indicating a map fragmentation event (which occur at upward threshold crossings). **C** A snapshot of the current observation (*z_t_*), which here consists of idiothetically centered BVC inputs to the agent (located at blue dot). Each pixel represents a BVC tuned to a location in space specified by the vector displacement between the agent and the pixel. Pixel intensity indicates the level of BVC activation. The image is cropped to exclude inactive BVCs. **D** Activity of the cell population encoding the STM (*m_t_*) of recent observations for the trajectory from A. The STM slightly lags the current observation. Our prediction signal is derived from a normalized dot-product between the current observation and the STM. **E:** At each time *t*, the model stores associations between the STM *m_t_*, the internal positional code *x_t_*, and its predecessor *x*_*t*-1_ in a long-term memory (LTM). The slots *s_t_* in the memory are chosen randomly. **F:** At a fragmentation event, the current observation *z_t_* is compared to the LTM of observations *M_z_* (blue columns), while the current position *x_t_* is compared with the corresponding LTM of predecessor positions *M*_. These comparisons result in weights *w_z_* and *w_x_*, respectively, and a memory is probabilistically selected for reuse from the LTM in proportion to its weight. Additionally, with a fixed probability, a random new map is initiated.

Once a decision to fragment has been made the agent must make a second decision, about which map fragment to use next, for which it uses its LTM. The LTM consists of past associations between the grid cell-encoded position representations and the sensory observations, filtered through the STM. Thus each LTM entry represents a gist of the sequence of observation-position pairings over an interval given by the time-constant of the STM (Fig. 3f). At the fragmentation event, the agent searches the LTM and stochastically selects a state in proportion to its match between items in LTM the current observation-position pairing. With some small constant probability (Fig. 3g), the agent selects (initializes) a new map, which corresponds to selecting a randomized new internal grid-coded position representation by randomizing the set of phases across the grid modules.

The stochastic selection of an item from LTM based on overlap with the current observation serves two purposes simultaneously: first, an observation is likely to drive selection of a closely matching prior observation, and second, the retrieval of a previous observation is also proportional to the number of times that observation has been made before, because stochastic selection from the set of past observations is a form of monte carlo volume estimation. In short, the selection of a submap after a fragmentation decision enables the reuse of existing submaps to represent new spaces when relevant based on similarity and frequency of past observations, while simultaneously permitting the creation of new maps. The frequency-dependence of this process together with the possibility to create new maps is similar to the Bayesian nonparametric Chinese Restaurant Process (CRP) [18, 30, 31]; the observational similarity component makes it more akin to the distance dependent CRP (dd-CRP) [32, 33]. However, in contrast with the CRP, observations in the present version of our model are only implicitly clustered into submaps: temporally-averaged observation-location pairs are stored independently of the rest in the LTM without an explicit submap assignment, with submap boundaries defined by the existence of a discrete fragmentation decision and a discontinuous jump in the grid-encoded spatial locations for the post-fragmentation observation relative to the immediate pre-fragmentation observation ^3^.

Further maintaining a “temporal” LTM which memorizes spatial transition probabilities, and using this information to bias the selection of a map at fragmentation events stabilizes how an environment is fragmented, though it is not critical (see SI, Fig. S6). Spatial transitions contain valuable information about the relationships between individual map fragments and are important for exploiting their hierarchical structure in route planning, as we illustrate later.

### Fragmented maps in multiple environments

We explore the map fragmentations generated by our online model across organically shaped and previously experimentally tested structured multi-compartment environments, Fig. 4. The online model generates fragmentations at locations that correspond to observation bottlenecks, including at doorways or narrow openings and around the corner of sharp turns, Fig. 4a-c (top). Starting from the first trajectory through the space and across multiple trajectories, the remapping or fragmentation points and the selected maps are consistent, evidence of the robustness and reliability of online frag-mentation (Fig. 4a-c, bottom). In the two-room and hallway environment, the model generates a fragmentation in which the two rooms are each represented by the same local map (rather than a single global map), and these maps are distinct from the map for the hallway. Moreover, the fragmentations generated by the online model are consistent with the fragments from the principled baseline method (Fig. 4d compared to Fig. 2e). Because of the stochastic process of map matching and retrieval from memory, new or multiple maps are sometimes formed and retrieved within in the same room (scattered gray dots not contained within highlighted gray regions, Fig. 4a-c, bottom). This suggests that there might be more than one map for the same space, even without contextual changes. This model can be used to generate fragmentations and predicted grid cell tuning curves for arbitrary enviromental geometries; we do so for model cells from different grid modules in two environments, Fig. 4e,h (fragmentations of more environments, including a square sprial maze and a simple linear track, shown in SI, Fig. S1a-c). If the angular field is of view *ϕ* is restricted rather than omnidirectional, the maps also acquire direction tuning, Fig. 4h and Fig. S1c.

**Figure 4:**
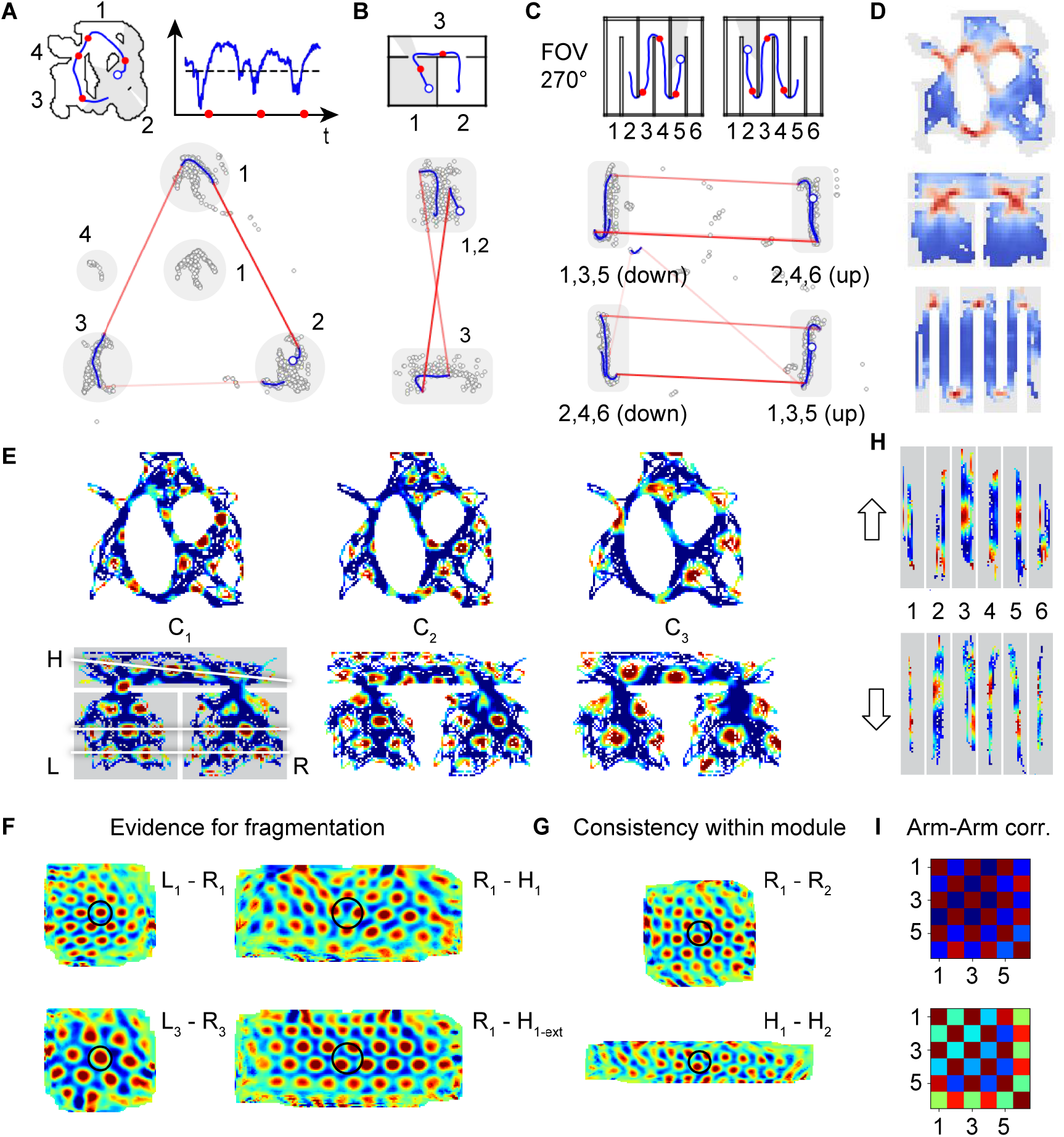
Online fragmentation results. **A–C** *Top:* Example trajectory snippets through three different environments (blue line), with current agent position indicated as an open blue circle. Fragmentation events indicated by red dots. The numbers indicate subregions identified in Figure 2E. A additionally shows the corresponding predictibility signal (blue) and threshold (dashed line). *Bottom:* The same trajectory snippets traversing the internal coding state space. The gray circular areas highlights the most-visited parts of state space for each environment, and the numbers correspond to the mapped area in the environment. Discontinuous jumps in state space, corresponding to transitions between submaps which occur at fragmentation events, are plotted in red. **D:** Heat maps indicating the density of online fragmentation events closely match the offline predictability-based clustering fragmentations from Fig. Figure 2E. The fragmentation decision map can be interpreted as the tuning curve of a “fracture” cell, which in multi-room environments can be interpreted as a “doorway cell” because the fractures happen at doorways. **E:** Firing fields of three simulated grid cells in two distinct environments. C1 and C2 are from a common module; C3 is from a distinct module of larger scale. White lines in bottom left panel show the submap discontinuity between the rooms and hallway. **F:** Evidence for map re-use and fragmentation: *Left:* Spatial cross-correlation of the left and right room representations (by C1 and C3) in the two-room-hallway environment is centered at zero (center of black circle), illustrating that the same map is reused in both rooms across modules. *Right:* Spatial crosscorrelation of the right room and hallway representations by C1 (top) and by a control pattern (bottom) that extends to the whole environment (in a continuous way without realignment) C1’s right room tuning curve. **G:** Cross-correlation of comodular cells (C1, C2) in the right room, and in the hallway: the cells have the same relative phase in each map fragment, showing maintained comodular cell-cell relationships within and across all map fragments. **H,I:** Directionally tuned firing fields in the hairpin maze (shown in C) of an idealized grid cell. The difference in firing fields for consecutive arms shows that the arms are mapped to different parts in mapping space depending on the direction they are traversed. The matrices show the correlation coefficients comparing signals of different arms.

### Coherence of fragmentation across scales and maintenance of cell-cell relationships

Two key structural predictions of our model are, first, that the map fragmentations are consistent and coherent across scales (across grid modules), with all cells and modules remapping at the same spatial location in an environment. This is in contrast with eigenvector-based models [20, 21, 34], in which there is no specific or coherent remapping decision that is made across eigenvectors, Fig. S3,S4, Fig. 6.

**Figure 5:**
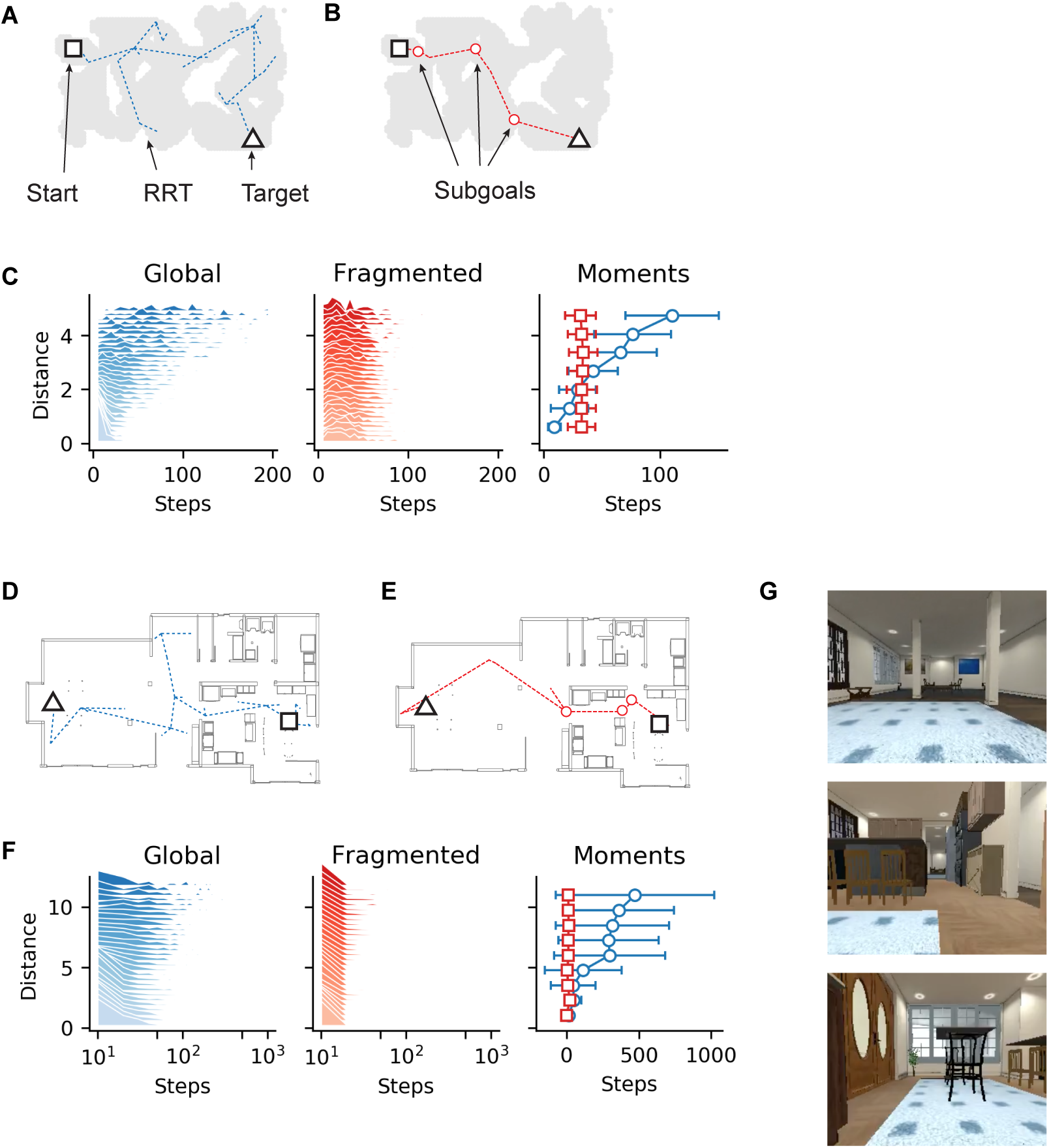
Efficient Hierarchical Planning with Map Fragmentations. **A-C:** Results of the planning algorithm applied to the global map of the environment (A) and to the hierarchical maps from our online fragmentation model (B). In C we plot the distributions of planning steps conditioned on the distance between start and target locations. **D-G:** Similar to A-C but using an alternative fragmentation algorithm based on semantic information extracted from visual inputs as shown in G.

**Figure 6:**
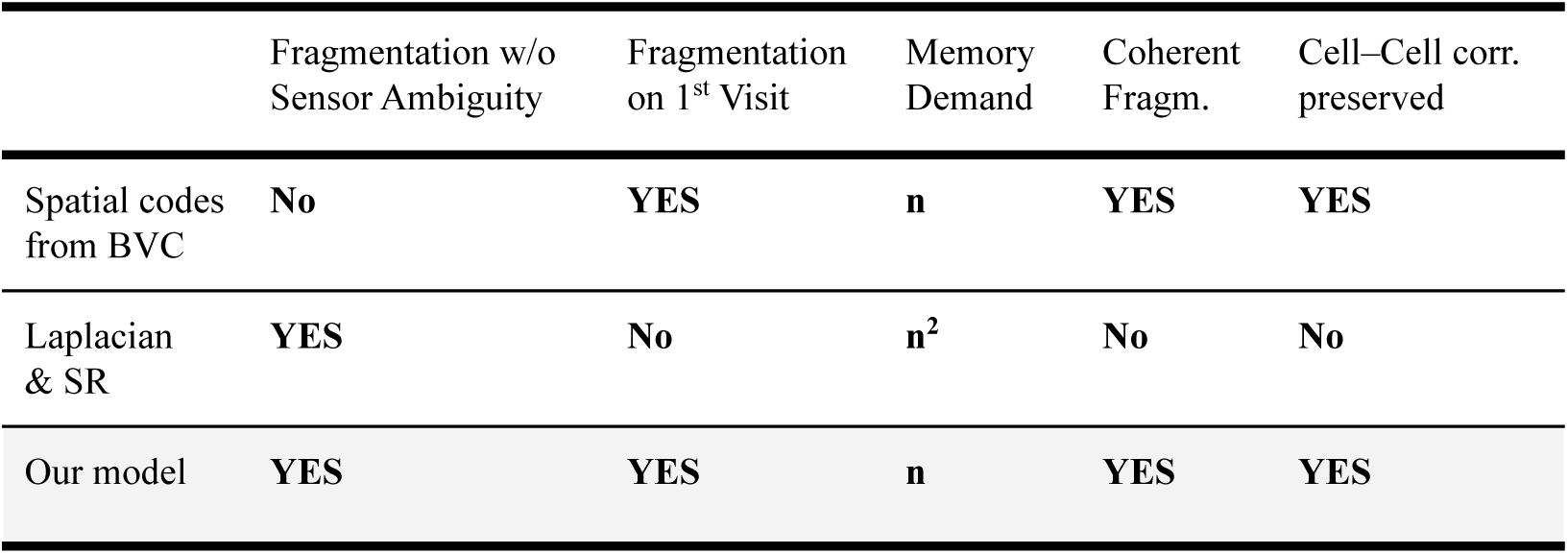
Comparison to other models. Our model improves on the potential shortcomings of other mapping approaches. Examples for model based on spatial codes derived from BVCs are [40, 41], and for models based on SR or graph Laplacian are [20, 21].

Second, in our model all grid cells within each module maintain fixed cell-cell relationships across map fragments and environments. This too is in direct contrast with eigenvector-based models, Fig. 4g,S5, Fig. 6. Consistent with our model, grid cell data and analyses reveal that the pairwise relationships between co-modular grid cells remain stable across environments [35] and states [36, 37] even when place cells remap and their relationships change. These neural data more generally do not support models in which grid cell responses are derived from place cell responses [34, 38] because they would predict altered cell-cell relationships when place cells remap [36].

### Efficient planning with fragmentations

Next, we quantify the functional utility of map fragmentation in a navigational planning problem. The fragmented maps, which represent a form of state abstraction, decompose the planning problem hierarchically, into a family of smaller and simpler sub-problems. Thus, they are expected to make planning more efficient. We perform computational experiments to illustrate this point, comparing a bi-level navigation algorithm in the fragmented map with a simple baseline.

Consider a goal-directed problem in which agents, who have previously mapped the space, are tasked with finding a path to a cued goal location from a start location. For planning, we will assume that the LTM containing stored observation-location associations also includes an explicit submap identification (that is, all observations until a fragmentation event are assigned the same submap ID; at a fragmentation event, if the retrieved map has not yet been assigned a submap ID, a new submap ID is initiated and added to the LTM and associated with all subsequent observations until the next fragmentation event, and so on; all the observations between fragmentations are fused using local displacement information to form a submap for the whole fragment) and storing submap transitions. The environments are complex, but *without* repeating submap structure (Fig. 5a-b, d-e), because the fragmented representations generated by our simple agent do not distinguish between difference spaces with the same appearance (no global odometry assumed across submaps).

The baseline (global) agent is furnished with a global map, which includes ground-truth position informaton for all observations (Fig. 5a,d) and uses the Rapidly-exploring Random Tree (RRT) algorithm [39] to find a path through the space (Methods). The agent using a fragmented approach constructs a graph in which the nodes correspond to the submaps, and the edges correspond to observed transitions between submaps during exploration. It performs a depth-first search through the transition tree to find the sequence of submaps that lead to the node containing the target location (determined by querying the LTM with the target inputs). Within each submap, the agent uses the RRT algorithm to plan a path between the locations corresponding to the entry and exit edges. This agent possesses no global positional information.

In the environment of Fig. 5a-b, routes are found vastly more rapidly with fragmented maps than without: we see a ~ 5-fold speedup. The relative advantage of planning with fragmented maps grows superlinearly with the complexity and size of the environment and separation between start and end locations within these spaces, Fig. 5c (right; steps are a proxy for the problem complexity).

Next, we simulate agents moving through 3D photorealistic virtual apartments in which observations are rich pixel images with range data, Fig. 5d, g. We apply convolutional visual recognition networks to the dense inputs to extract sparse landmarks and use these to generate online map segmentations (Methods). As before, the agent performs bi-level planning on the tree of transitions with submaps as nodes and RRT planning within submaps. Here we find a several orders of magnitude speedup in planning with map fragmentation, Fig. 5f.

## Discussion

### Relationship to existing work

Existing models of neural representations in multi-chambered environments fall into three categories, Fig. 6: In the first, fragmentation is driven by path integration errors that cause a large mismatch between estimated position and familiar observations [40]; in environments with little ambiguity in the external sensory cues or no path integration errors, there would be no fragmentation. In the second, spatial representations are assumed to be derived directly and entirely from combinations of external cues, thus similar external inputs induce similar representations, but without a notion of map fracturing or explicit discrete fragmentations [41]. The third category represents positions in an environment as states, and spatial representations as the top eigenvectors of the transition matrix between states (the transition matrix can be defined by how a specific agent or random agent traverses the states) [20, 21]. These models require a global and veridical acquisition of a complete map of the environment and of the transition matrix between all pairs of locations in the environment, before any potential spatial representations can be defined. As in the second category, these also do not provide explicit or discrete fragmentations of the environment; different eigenvectors have different spatial patterns, changing at different spatial frequencies across the space. If two eigenvectors with the same spatial frequency are interpreted as two co-modular grid cells, they do not maintain their cell-cell relationships across the space, in contrast with the known preservation of comodular grid cell phase relationships not just within but across environments, time, and behavioral states like waking and sleep [35–37, 42]. By contrast, our model is fully online so that from the very first trajectory in an environment it generates explicit, robust fragmentations of the space at specific discrete locations in each space. It does so by driving a discontinuous and synchronized randomized phase change across all grid modules at the fracture point. These fracture events are determined by a rise in surprisal or prediction error. This induces fractures across regions whose geometries are quite distinct, e.g. a doorway, a u-turn in the hairpin maze, or a narrowed hall-like passage in the organically shaped environment (cf. Fig. 4). The fragmentations are predicted to occur even in the absence of positional ambiguity from path integration error, preserve co-modular phase relationships across all map fragments, and require a much smaller time and memory demand than transition matrix models. Finally, the model involves only simple, biologically plausible computational elements, with grid cells and BVCs, a short-term memory, and a long-term memory, to explain a number of experimental results.

Our work, initially motivated by the empirical observations of fragmented maps in neuroscience, is closely related to work on segmented maps in the field of simultaneous localization and mapping (SLAM) in robotics [19, 43–45]. The main difference is that predictions in our model are based on a temporally limited window into the past, provided through the STM, whereas in [19] *all* observations are accumulated into a map that the prediction is based on. Further, our predictions are based on idiothetically-centered local views of the environment (BVC) – which are not assembled into a global allocentric map – and use an adapted moving average as a STM. For (re-)localization we use local views stored in a spatially indexed LTM.

As we have shown, spatial abstraction and spatial hierarchy in the form of map fragments can be of high utility in efficient search for solving goal-directed problems. State abstractions and hierarchical representations are broadly recognized to be important for more efficient reinforcement learning as well, and implemented in different forms including the classic options framework and more recent attempts [46, 47]. A key challenge for such approaches is to find rules that generate appropriate state abstractions, especially those capable of doing so in an online or streaming way. Our work is a contribution in this direction; related work includes the generation of temporal abstractions based on novelty rather than surprisal [48].

Our use of a surprisal signal is closely related to curiosity-based algorithms for reinforcement learning [49]. These algorithms use prediction error as an internal reward, to drive agents to explore unknown parts of the space. By contrast, we use prediction error as a way to generate state abstractions.

### Model extensions: a broader set of metrics for fragmentation

The general principle of online state abstraction through online map fragmentation can use metrics in addition to surprisal for triggering a fragmentation event. Consider the case of two hallways with similar ideothetically-centered views, e.g. hallways 2 and 3 in Fig. 4c, that differ only in the permitted turn direction at the end. A natural extension of the model would be to incorporate a cell population encoding navigational affordances, to fragment and select maps based not only sensory surprisal but also on the set of actions that can be or are commonly taken. Other extensions include using the physical distance between states [20, 21], the passage of time [50–52] with a dynamic (temporally decaying) threshold for fragmentation that makes fragmentation more likely as time elapses (also see [19]), the appearance unique or novel visual features including landmarks [48, 53–55], and sufficient mismatch in the estimates of state made from different cues or sensory modalities [40, 56], in addition to the metric of perceptual predictibility that we have used here and that the hippocampus has been shown to be sensitive to [57, 58]. The present model, which provides an online method for generating meaningful abstractions, may be applied with arbitrary combinations of these metrics to generate fragmentations influenced by multiple factors.

### Merging of maps

In case of prolonged experience in the two compartment environment, map fragmentations tend to merge into a single, continuous representation that covers both compartments [14]. In our model some map fragments can, because of the stochastic nature of the fragmentation process, occasionally extend beyond an expected fragmentation boundary (see Fig. S1d). These events occur sparsely and are unlikely to be the source of the merging of maps observed in [14]. We expect the merging of maps to result from an improvement of the prediction signal with more experience, which can be modeled by allowing the prediction system to use not just recent observations from short term memory, but also past observations from long term memory. Exploring the dynamics of this process is an interesting potential extension of the present work.

### Role of map fragmentation for general cognitive representation

Our model of online fragmentation of a continuous stream of experience enables the representation of a very general class of maps – including in spatial and non-spatial cognitive domains – in a way that exceeds the capabilities of a “pure” grid code. Grid cells generate Euclidean representations of Euclidean spaces [11]. Fragmented maps can each be viewed as separate local Euclidean “charts”, mapped out by a multi-modular grid code, that are then associated to each other through transitions learned in the hippocampus according to the global layout of the charts. In other words, the combination of fragmented maps and the transitions between them can be viewed as a topological atlas [59] or topometric map [60], that can represent highly non-Euclidean structures while also permitting locally metric computations.

Thus, from a general perspective, map fragmentation and remapping (reanchoring) on cognitive representations can be viewed as faciliating the step from representing flat Euclidean space to representing richer manifolds. In combination with grid cells’ ability to represent high-dimensional variables [11], such a coding scheme becomes highly expressive.

In contrast to the approach taken in [12, 61] there is no need to generate entirely new neural codes and representations to fit the local statistics of ea explored space. Instead, we propose that the neural codes seen within submaps retain their native structure across spaces, in the form of a pre-formed and stable recurrent scaffold for memory through grid cells. Even though grid cell representations in each module are 2-dimensional, theoretically the set of modules an represent even high-dimensional continuous spaces [11], while potential non-Euclidean aspects of cognitive varaibles can be captured by the between-submap transitions. This structuring of memory into continuous parts with preexisting scaffolds [62–66] together with occasional transitions between these continuous chunks simultaneously provides rapid learning and flexibility.

### Episodic memory

Episodic memory, one example of a general cognitive representation, deserves special dis-cussion because of the privileged role of the hippocampal system in its creation, storage, and use [67, 68]. Like spatial map fragmentations, episodic memory involves fracturing the continuous stream of temporal experience into chunks that involve similar perceptual, temporal, and contextual elements [3–5, 69]. Thus, our proposal for surprisal- or prediction-error based spatial segmentation can, with minimal modifications, be applied to study memory chunking. Interestingly, the memory for non-spatial items has also been shown to segment based on changes in spatial context, specifically by passage through doorways [70], as would be predicted by the present model.

The utility of applying our model first in the spatial domain is that it yielded concrete predictions that we found to be quantifiably consistent with observed neural representations and map fragmentations. Applying it next across cognitive domains will contribute to a unified computational model for how the hippocampal formation generates structured memories of spatial and non-spatial cognitive experience [12, 18, 61, 68, 69, 71, 72], and how these fragmented representations could permit more efficient and flexible use of memory for cognitive problem solving.

### Experimental Predictions

The decision to form a new map fragment in our model depends only on recent observations that are filtered through a STM, without requiring global information about the enironment. Thus, map fragmentations are predicted to occur in real time and on the very first pass through relevant regions of new environments, consistent with experimental results in the spatial and non-spatial domains [3–5, 73]. Further, in our model, all grid cells and grid modules undergo map fragmentation simultaneously, at the same time and location along a given trajectory, unlike in other models (Fig. S3,S4) [20, 21].

Fragmentations tend to occur at spatial bottlenecks that limit the prediction horizon, which correspond to “doorways” in the environment. The current evidence for cells firing at doorways is mixed [74, 75]. However, the necessity for a neural correlate that communicates the fragmentation decision and facilitates across-module grid realignment under a fragmentation event predicts the existence of “fracture cells” whose tuning curves would resemble the heatmaps of Fig. 4d, which in these environments are consistent with an interpretation as “doorway cells”. Note that regions at which a fracture cell would be active can correspond to locations with quite dissimilar local geometries, as seen in the fracture locations across three distinct contexts in Fig. 4d; thus, fracture cell tuning curves are not merely an instance of place field repetition caused by similar local geometries [41].

A common theme in MEC seems to be that cells with spatially structured tuning coexist with vector versions of themselves: i.e., cells that have similar tuning curves but are offset by a fixed vector (e.g. BVCs [26, 27] and landmark or object vector cells [76, 77]). In this light we might also expect “fragmentation vector cells” or “doorway vector cells” that fire if the rodent is at a fixed angle and distance from a fragmentation location. These cells, which could be interpreted as encoding future action affordances or future map transitions, would faciliate planning.

Next, the model predicts that the stochastic process of generating map fragmentations can result in more than one map for the same region even when there is not an explicit manipulation of context or task. There are at least two implications of this result. First, it suggests that variations in the firing of grid and place cells on different visits to a location might be due not only to variable paths taken within a single map [78] but to the retrieval of entirely different maps. Second, these multiple, stochastically generated maps might subsequently be easy to harness for contextual differentiation, for instance like “splitter” cells [61, 79–81].

Finally, the large efficiencies in planning and goal-directed navigation afforded by the use of fragmented maps suggest that neural planning should exhibit hallmarks of the fragmentation process: If theta phase precession or waking neural replay events [82–88] correspond to planning [89–91], we should expect them to exhibit punctate trajectories with hierarchical dynamics between versus within fragments in multicompartment environments.

## Methods

### Data and code availability

The source code will be made available on request and posted online upon publication.

### Offline fragmentations from predictability and surprise

We approximate the probabilistic observation model *P*(*z’* | *x’,x,z*) by

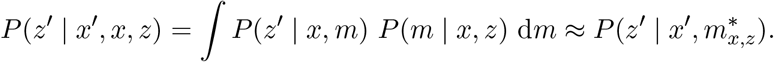

Here 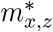 is the maximum a posteriori estimation of an occupancy map given by an inverse sensor model as described in [29], and *P*(*z’* | *x, m*) is the respective range sensor model. More precisely: Given a deterministic range sensor that takes measurements along a fixed number (*n* = 1000, 1500) of simulated beams, whose angles are chosen at equally spaced angles from the interval [−*π*, *π*], we take three depth measurements *z*, *z’*, and *z’’*. The first two are taken in the actual environment at their resepective poses *x* and *x’*, whereas the third is taken on a map 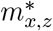 built from the initial measurement *z* made at *x*. The observation model *P*(*z’* | *x, m*) is then defined as a multivariate diagonal Gaussian with constant diagonal entries *σ* = 1.0 and mean *z’’*, Fig. 2a,b.

The function underlying the distance matrix used for the Isomap embedding (cf. Fig. 2d) is given by the *mutual surprise s*(*x*, *x’*) between two poses *x*, *x’* which we define as

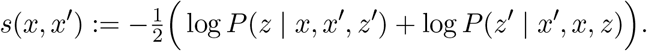

We refer to the negative mutual suprise as *mutual predictability*. With this in hand we define the *average surprise s*(*x*) = *s*(*x*; *r*, *ε*) of a pose by averaging over the mutual surprise about all poses at a fixed distance. To be more precise we define

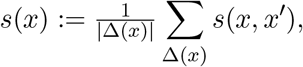

where Δ_x_ = Δ_x_(*r*, *ε*) is the set of all poses whose distance to *x* lies in the range [*r*–*ε*, *r*+*ε*], for some previously fixed *r*, *ε* > 0; in our experiments we use *r* ≈ 0.4 and *ε* ≈ 0.05 depending of the minimal distances between poses. We sometimes refer to the negative avergage mutual suprise as *contiguity*. Informally, a high contiguity implies fewer suprises in direct proximity of the current pose and thus a low urge to remap.

To extract map fragmentations we uniformly sample poses from the environment and compute their avergage surprise, Fig. 2e. We then consider only those poses whose surprise lies below a previously fixed threshold (chosen acordingly for each environment). To make an informed choice about the threshold we compute a discrete contour tree [24] of the poses with respect to the average surprise visualizing the evolution of the connectivity with respect to increasing thresholds, Fig. 2f. The connected components of the subthreshold region yields a fragmentation into sub-maps, one for each connected region, and a suprathreshold transition region. We consider two poses to be connected if their Euclidean distance is below another previously fixed threshold that depends on the coverage of the environment by all the pose samples.

### Online fragmentations from predictability

#### Observations and internal mapping locations

As before, our observation model is given by a range sensor that takes measurements along a fixed number of simulated beams. The beams’ angles are chosen at equally spaced angles from the interval [*θ_t_* – *ϕ*/2, *θ_t_* + *ϕ*/2]. Here *θ_t_* denotes the head direction at time *t* and *ϕ* = 360°, 270° defines the field of view of the agent; cf. Fig. 3a. We convert these range measurements into the activity *z_t_* of a simulated population of boundary vector cells by a binning process; cf. Fig. 3a,c. In our model we assume there is a *n* × *n* array of BVCs covering an area of *w* × *w*, with *n* = 91, 111 and *w* ≈ 4m.

We assume that internally locations are represented by a population of idealized grid cells of mulitple scales. For ease of computation, we interpret this multi-module grid code as a high capcity code for an unfolded 2-dimensional space [11]; cf. Fig. 4a-c (bottom). The Poisson rate maps *f_c_* for an idealized grid cell *c* are then generated from superposing three cosinusoidal waves, each offset by an angle of 60^°^, over the unfolded 2-dimensional grid space, i.e.

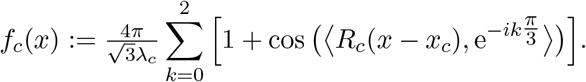

Here *λ_c_* and *x_c_* encode the lattice scale and its offset, and *R_c_* is a rotation matrix defining the orientation of the lattice.

#### Short term memory

The short term memory (STM) is defined as an adapted exponential moving average of BVC activity:

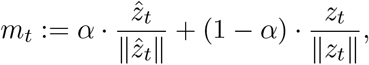

where the prediction

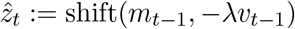

is a shifted version of the 2d-array m_t-1_ with respect to the scaled velocity of the agent. We found that a smoothing parameter *α* ≈ 0.9 works well. The scaling parameter 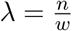 maps from the environment into pixel space. The shift of the BVC array results in a diffused version of the array caused by shifts with non-integer values. The extent of diffusion depends on the resolution (or number) of the BVCs.

#### Prediction model and fragmentation events

The prediction model is a normalized dot product of the current BVC observation *z_t_* with the prediction *ẑ_t_* computed from the STM as described above:

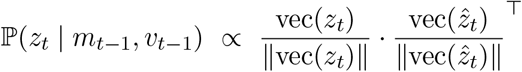

where vec(*z*) is the unfolded version of a 2d-array *z*; cf. Fig. 3b-d. A fragmentation event is triggered after the prediction signal 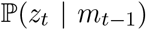 recovers from falling below a fixed threshold *θ* (≈ 0.9, 0.925) and rises above again; cf. Fig. 3b. The normalization and the fact that both *z_t_* and *ẑ_t_* are non-negative ensures that the prediction score lies within the intervall [0, 1].

#### Long term memory and relocalization

The LTM is implemented as a matrix 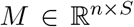 whose *s*’th column is given by the concatenation of the internal position estimate *x_t_s__*, its predecessor *x_t_s__*–1 and the state *m_t_s__* of the STM at the time *t_s_* the entry was written to memory, i.e. we have (cf. Fig. 3e)

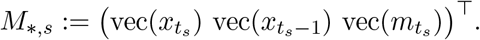

We fill the memory as follows: At each time step *t* we choose a slot (column) *s_t_* in the memory and replace the corresponding entry with the new one. Until we reach capacity, that is as long as ≤ *S*, we set *s_t_* = *t*, after that the slots *s_t_* are chosen uniformly at random – similar to the associative memory in [92]. Thus, the LTM consists of two associative memories: one storing assiciations between locations and observations, and the other storing state transitions. Alternatively, one could store associations with the actual observations *z_t_s__* and not the filtered observations from the STM *m_t_s__*, but we found that the associations with the STM work better and result in more stable fragmentations. The same is true when we restrict the capacity of the memory; cf. Fig. S6. Note that the LTM also maintains a temporal memory storing transitions (*x_t_s_–1_*, *x_t_s__*) for each entry in the memory. We use a memory size *S* between 2000 and 6000.

To determine the new location during a fragmentation event we query the LTM and compute two distinct weight vectors *w*_1_ and *w*_2_. The first encodes how well a given observation *z* fits any of its entries and is given by

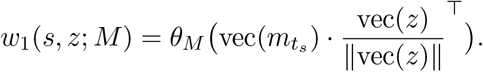

With slight abuse of notation we denote by *θ_M_* the function that sets all values below a certain threshold *θ_M_* to −∞. For ease of notation we set e^−∞^:= 0 – this becomes relevant in the probability computation below. We usually set this threshold to be equal to the fragmentation threshold *θ* ≈ 0.93. In order to allow for more flexibility during the above lookup we query the LTM not only with the actual observation *z*, but also with observations shifted by small pixel offsets *δ*, i.e. with *z_δ_* = shift(*z*, *δ*) instead of just *z*, where 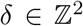 is chosen from a small region Δ around the origin. If a shifted observation fits a particular entry in the memory better, we replace the corresponding entry in the computed weight vector *w*_1_. Then, if one of these adjusted entries, *s* say, is chosen during a remapping event we do not remap exactly to the associated position *x_t_s__* but adjust it proportional to the respective offset *δ* and remap to 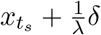 (recall that λ translates from environment to pixel coordinates).

The second vector serves as a bias towards map transitions that have already been traversed and is given by

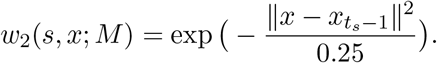

Note that we use the Euclidean norm between two 2d vectors out of computational convenience, but we could have used the dot product of their corresponding multi modular grid codes as well. Finally, when a fragmentation event is triggered we sample a new location from

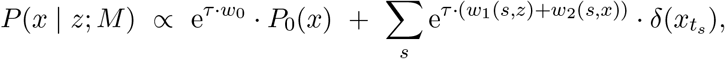

where *P_0_*(*x*) is a distribution over the space of possible locations, *w*_0_ = 1 the concentration parameter, and *τ* = 1, 10, 20 is the inverse temperature of the model; cf. Fig. 3g.

#### Trajectories

The trajectories are generated by choosing waypoints in the environment uniformly at random and navigating toward the next waypoint along a perturbed shortest path at a mean speed of 20 cm/sec. Time is discretized into steps of size Δ*t* = 0.1 sec.

#### Hierarchical Planning

We apply Rapidly-exploring Random Trees (RRT) [39] to find a path between randomly chosen pairs of start and target positions, Fig. 5a. Next, we run our online segementation algorithm to get an environment-fragmentation into submaps and form a topological graph whose vertices and edges correspond to submaps and transitions respectively. For each map-fragment we superpose all its associated memories (STM-filtered BVC activity) and threshold this newly formed representation to form an occupancy grid map (in the sense of [29]) which we can apply the path planning algorithm to. We exploit the hierarchical structure by first finding a path of transitions in the topological graph, using a breadth-first search, and reduce the overall planning task into a family of sub-problems as follows: Each transition into- and out of a node defines a pair of local entry and exit postions on the submap associated with the traversed node defining a smaller planning problem that can be solved more effectively, Fig. 5b. In Fig. 5c we plot the distances between start and goal locations against the number of planning steps needed.

The algorithm underlying the results in Fig. 2d–g is given as follows. Because the 3D environments involve dense observations of pixel-rich data, we add image processing and observation sparsification steps in the form of landmark identification. The agent receives RGB-D images as input, removes the floor plane, and segments the resulting point cloud. It retains as landmarks the large segments that are not vertical walls, which are generally large furniture items that are both relatively static and easy to recognize robustly from new viewpoints. As it moves through the environment, fragments are defined as follows: Starting at the initial location, the current fragment is defined a set of two visible landmarks, and the region of space from which both those landmarks remain in view constitutes the set of spatial locations assigned to that fragment. Whenever the agent moves into a part of the space where one or both of those landmarks are not visible, and if the current location does not correspond to any existing fragment, it starts a new fragment. Each fragment is connected topologically to the fragment it entered from.

#### Arm-arm correlation

The correlation matrices in Fig. 4i are computed as follows. For each arm in Fig. 4h we produce a 1-dimensional signal by averaging over the x-axis of the respective tuning curves in each arm. Each entry *c_ij_* in the matrix is then given by the Pearson correlation coefficient of the 1-dimensional signals in arm *i* and *j*.

## Footnotes

1 The density notion in DBSCAN is based on a count of neighbors. We use the average mutual surprise instead.

2 This is also known as a grid occupancy map in robotics [29].

3 A temperature hyperparameter controls the degree of noise in the selection of a submap from LTM. This stochastic process allows us to not only use the degree of similarity but also the frequency of similar observations in selecting a submap: it performs a stochastic measurement of the volume of similar observations (submap occupancy), and then stochastically selects a map on that basis, without keeping an explicit count of how often each submap has been visited in the past. Thus, we may call this process a doubly-stochastic dd-CRP.

