## Supplemental Figures for "Fragmented Spatial Maps from Surprisal: State Abstraction and Efficient Planning" (from another version of this preprint: version 1, 2021-11-01)

**List of supplementary figures:**

1. Addtional Results.
2. Additional Firing Fields and Autocorrelations.
3. Eigenvectors of the SR (Organically shaped environment).
4. Eigenvectors of the SR (Two-room environment).
5. Within-Module Cross-correlation: SR vs. Our Model.
6. Effect of Memory Capacity on Map Number.

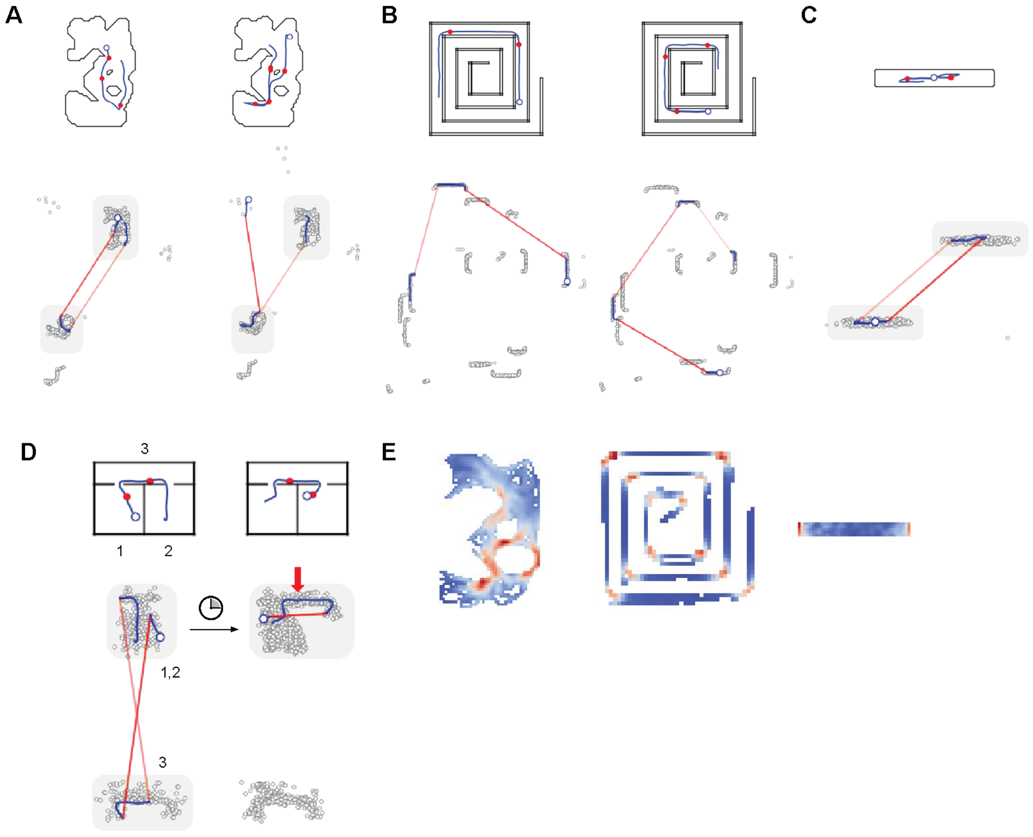


**Figure S1: Addtional Fragmentations.** **A-C:** Additional trajectories for three different environments (top) and their corresponding counterparts in mapping space (bottom) are shown. In the 1D track, the agent's field of view is less than 360 degrees, which results in different map fragments for the two directions of travel in the track. **D:** An occurence where a map fragment extends beyond the expected submap boundary. The arrow points to an obscured fragmentation event where the newly chosen location is close to the old position before the fragmentation event. **E:** Surprisal maps for the environments in (A-C).

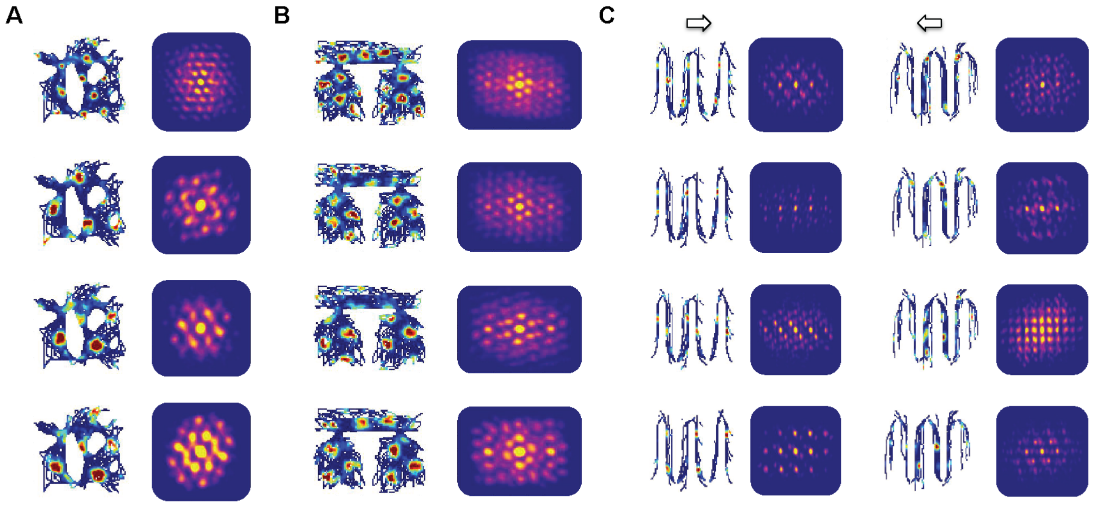


**Figure S2: Additional Firing Fields and Autocorrelations.**

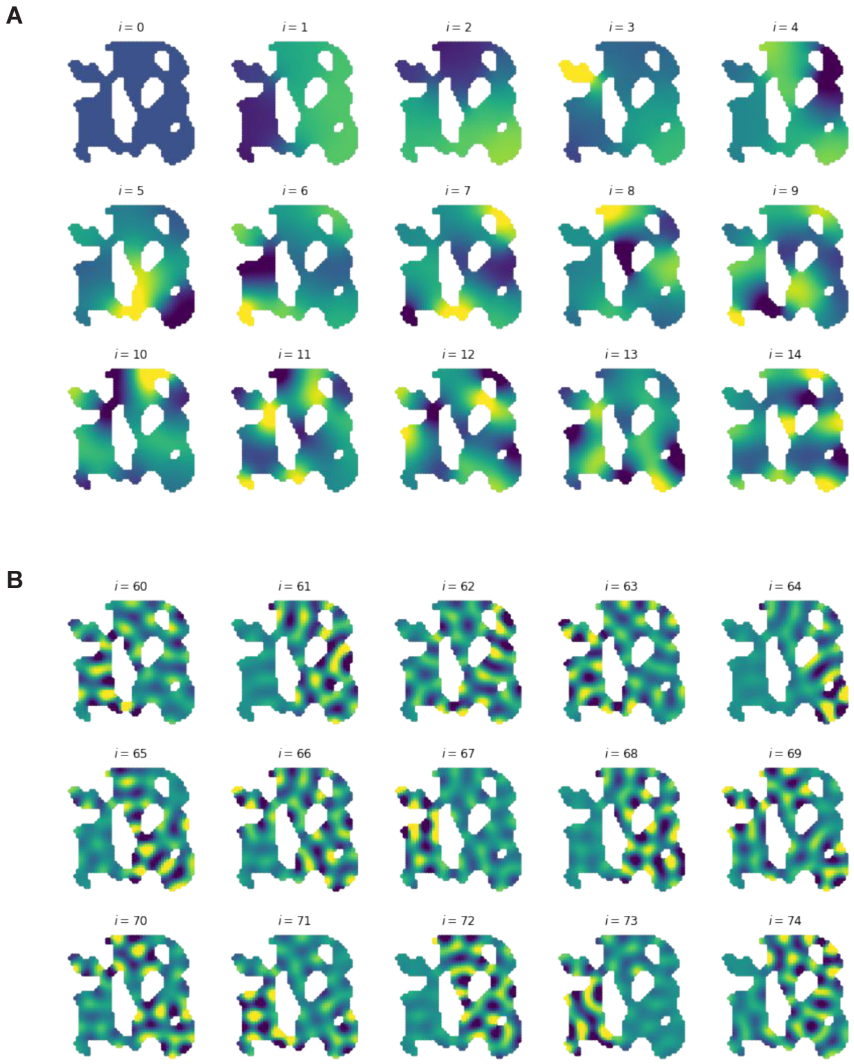


**Figure S3: Example Eigenvectors of the SR in our organically shaped environment, for comparison with our model's predictions.** Eigenvectors corresponding to the 1st-15th (A), and 61st-75th (B), largest Eigenvalues of the SR with $\gamma = 0.95$.

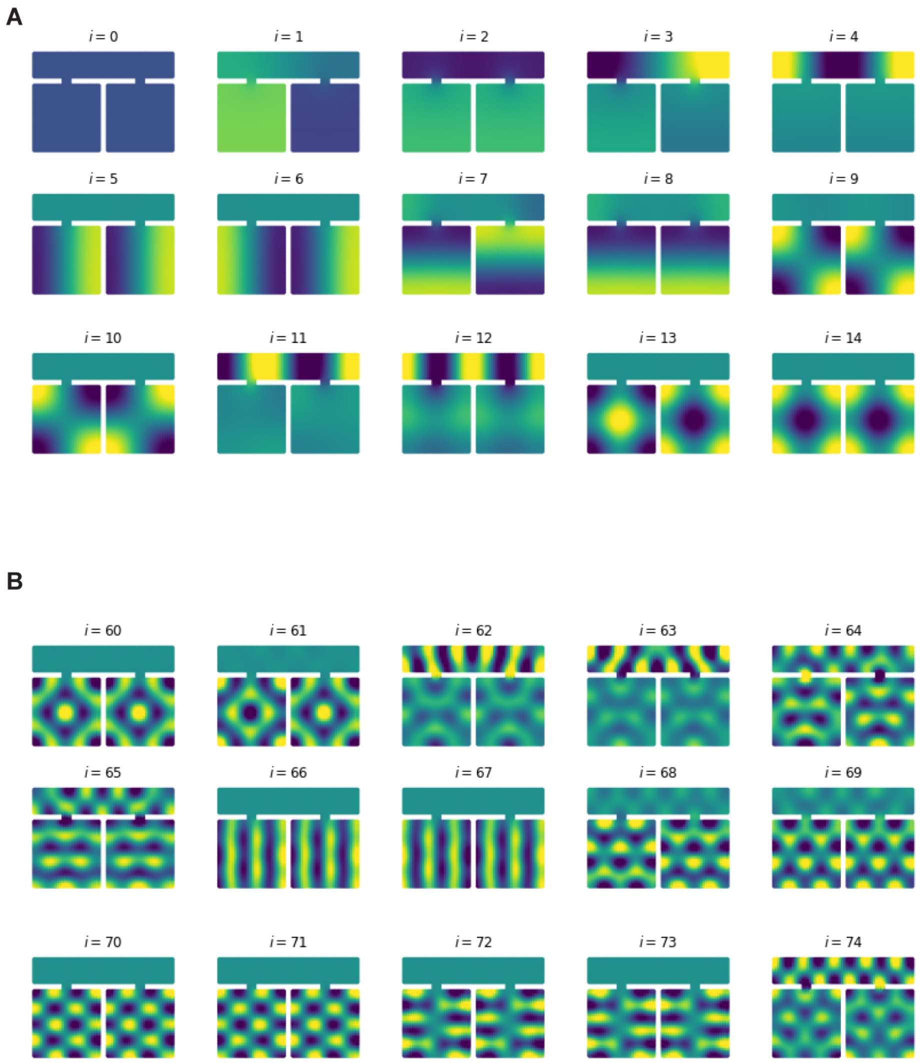


**Figure S4: Example Eigenvectors of the SR in the 2-room environment.** Eigenvectors corresponding to the 1st-15th (A), and 61st-75th (B), largest Eigenvalues of the SR with $\gamma = 0.95$. Note that SR "neurons" (eigenvectors) do not maintain a fixed relative phase difference in different parts of the environment, in contrast with empirical findings that show that grid cell phase relationships remain stable across time [1], environemnts [1, 2], and behavioral states including waking and sleep [3, 4].

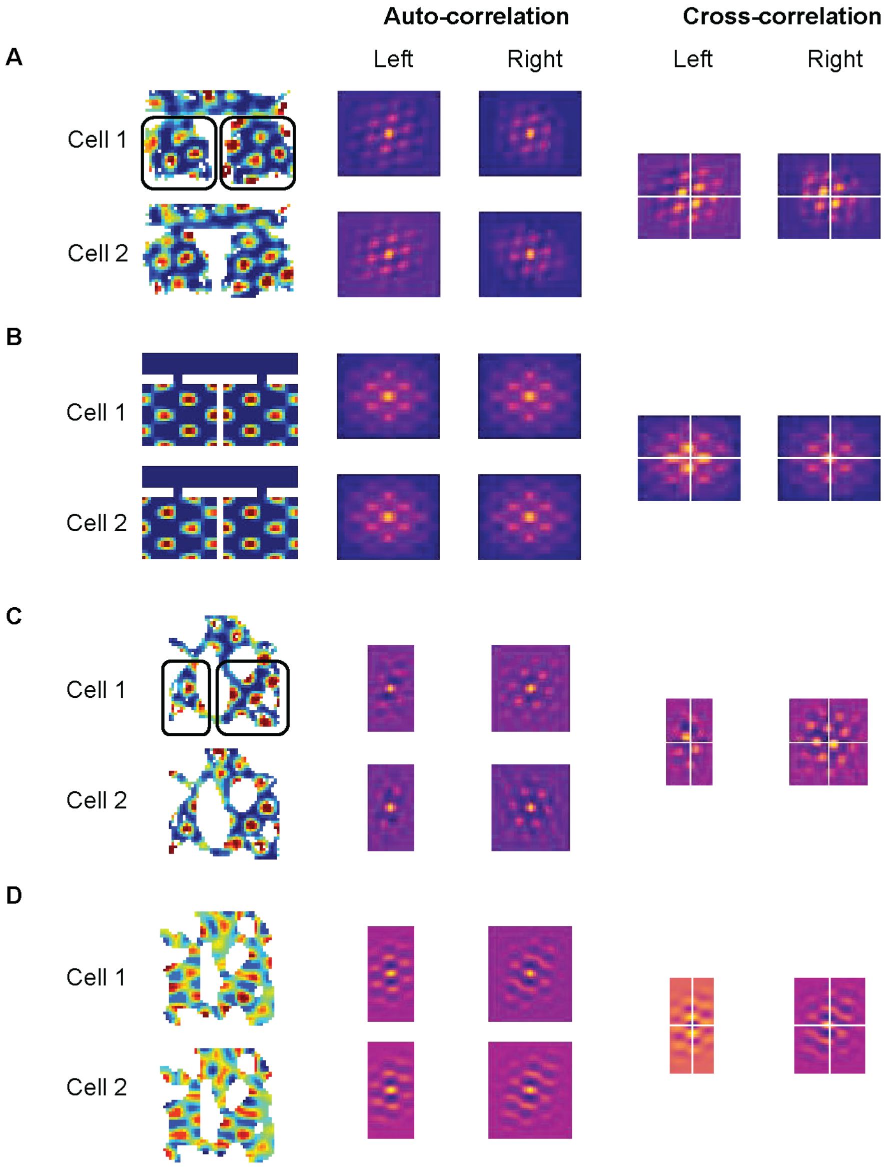


**Figure S5: Within-module cell-cell cross-correlations are preserved across map fragments: Our Model. A-D:** *Left, Middle.* Firing fields of two idealized grid cells from a common module and their autocorrelations in each of the highlighted compartments. In (B) and (D) the firing fields correspond to Eigenvectors of the SR of similiar scale. *Right.* The cross correlation of the two cells with each other in different parts of the environment: cross correlations and thus cell-cell phase relationships are preserved across map fragments (A,C), in contrast with the SR Eigenvectors (B,D).

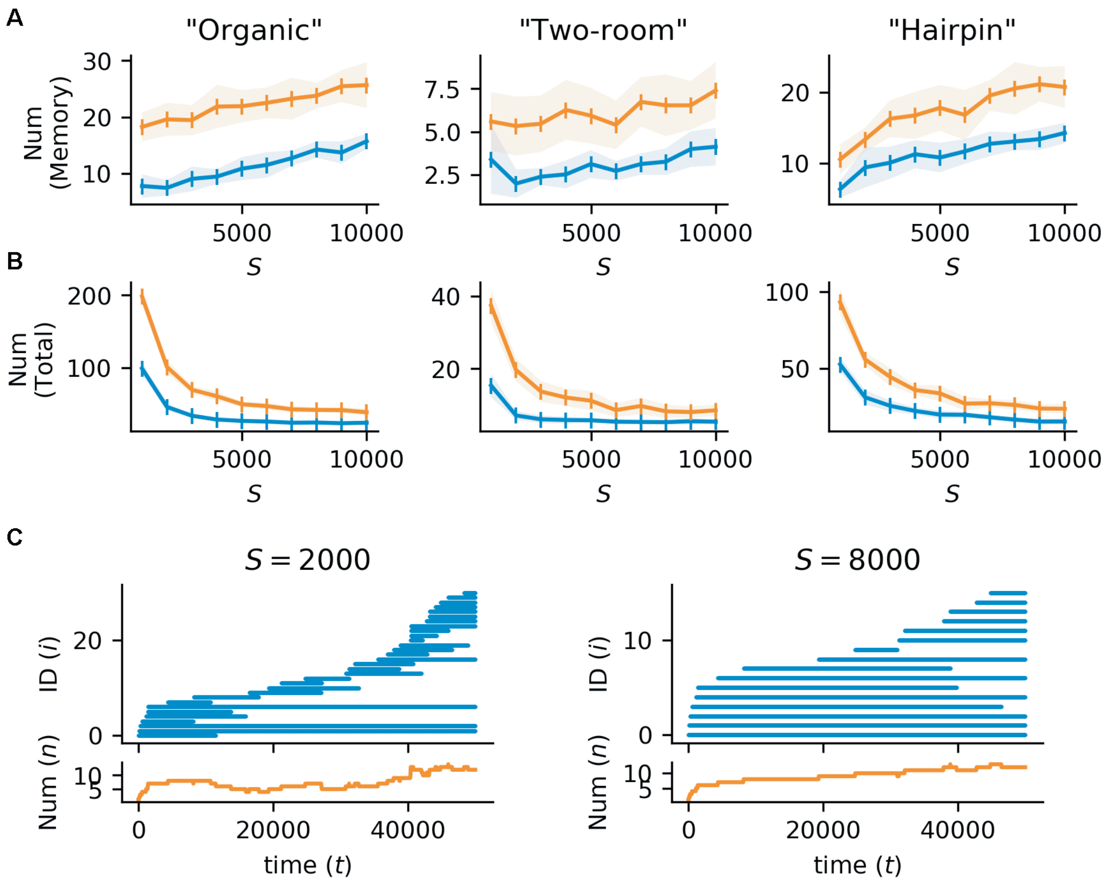


**Figure S6: Effect of Memory Capacity on Map Number. A:** We plot the mean number of maps in memory after 50,000 timesteps over a set of 15 trials. The plot compares trials where the temporal part of the LTM memorizing state transitions is switched off (orange) and where it is switched on (blue). The plots clearly show that the number of maps and the capacity of the memory are correlated. **B:** We plot the total number of maps started – and not just those maps stored in the memory. Interestingly the number of total maps is negatively correlated with memory capacity. **C:** The blue scatter plot indicates which maps are in memory at each time, plotting a marker at $t, i$ if map $i$ is in memory at time $t$. The orange plot shows the number of maps in memory at time $t$. We contrast two trials with low ($S = 2000$) and high ($S = 8000$) capacity.
